# Heritability of regional brain volumes in large-scale neuroimaging and genetic studies

**DOI:** 10.1101/208496

**Authors:** Bingxin Zhao, Joseph G. Ibrahim, Yun Li, Tengfei Li, Yue Wang, Yue Shan, Ziliang Zhu, Fan Zhou, Jingwen Zhang, Chao Huang, Huiling Liao, Liuqing Yang, Paul M. Thompson, Hongtu Zhu, Pediatric Imaging, Neurocognition and Genetics (PING), Connor McCabe, Linda Chang, Natacha Akshoomoff, Erik Newman, Thomas Ernst, Peter Van Zijl, Joshua Kuperman, Sarah Murray, Cinnamon Bloss, Mark Appelbaum, Anthony Gamst, Wesley Thompson, Hauke Bartsch, Alzheimer’s Disease Neuroimaging Initiative (ADNI), Michael Weiner, Paul Aisen, Ronald Petersen, Clifford R. Jack, William Jagust, John Q. Trojanowki, Arthur W. Toga, Laurel Beckett, Robert C. Green, Andrew J. Saykin, John Morris, Leslie M. Shaw, Zaven Khachaturian, Greg Sorensen, Maria Carrillo, Lew Kuller, Marc Raichle, Steven Paul, Peter Davies, Howard Fillit, Franz Hefti, Davie Holtzman, M. Marcel Mesulman, William Potter, Peter J. Snyder, Adam Schwartz, Tom Montine, Ronald G. Thomas, Michael Donohue, Sarah Walter, Devon Gessert, Tamie Sather, Gus Jiminez, Danielle Harvey, Matthew Bernstein, Nick Fox, Paul Thompson, Norbert Schuff, Charles DeCarli, Bret Borowski, Jeff Gunter, Matt Senjem, Prashanthi Vemuri, David Jones, Kejal Kantarci, Chad Ward, Robert A. Koeppe, Norm Foster, Eric M. Reiman, Kewei Chen, Chet Mathis, Susan Landau, Nigel J. Cairns, Erin Householder, Lisa Taylor-Reinwald, Virginia M.Y. Lee, Magdalena Korecka, Michal Figurski, Karen Crawford, Scott Neu, Tatiana M. Foroud, Steven Potkin, Li Shen, Kelley Faber, Sungeun Kim, Kwangsik Nho, Leon Thal, Richard Frank, Neil Buckholtz, Marilyn Albert, John Hsiao

## Abstract

Brain genetics is an active research area. The degree to which genetic variants impact variations in brain structure and function remains largely unknown. We examined the heritability of regional brain volumes (p ~ 100) captured by single-nucleotide polymorphisms (SNPs) in UK Biobank (n ~ 9000). We found that regional brain volumes are highly heritable in this study population. We observed omni-genic impact across the genome as well as enrichment of SNPs in active chromatin regions. Principal components derived from regional volume data are also highly heritable, but the amount of variance in brain volume explained by the component did not seem to be related to its heritability. Heritability estimates vary substantially across large-scale functional networks and brain regions. The variation in heritability across regions was not related to measurement reliability. Heritability estimates exhibit a symmetric pattern across left and right hemispheres and are consistent in females and males. Our main findings in UK Biobank are consistent with those in Alzheimers Disease Neuroimaging Initiative (*n* ~ 1100), Philadelphia Neurodevelopmental Cohort (*n* ~ 600), and Pediatric Imaging, Neurocognition, and Genetics (*n* ~ 500) datasets, with more stable estimates in UK Biobank.

---

The contribution of genetic variations to brain structure and function is of great interest. One major goal of brain imaging genetic studies is to understand the degree to which genetics can explain variations in imaging phenotypes, which are usually measured by the associated heritability. Heritability is the proportion of observed phenotypic variation that can be explained by the inherited genetic factors. By measuring the relative size of genetic and non-genetic effects on phenotypic variance, heritability can provide insight into the genetic basis of a phenotype and guide downstream analysis on more specific biological questions. Specifically, heritability can be measured by either the proportion of total genetic variation (broad sense), or the proportion of total additive genetic variation (narrow sense) (Visscher et al., 2008). One traditional way to estimate narrow-sense heritability is using samples from twin/family studies (Bartels et al., 2003; Visscher et al., 2006), in which the pedigree information can capture the effects of all genetic variants on phenotype (Visscher et al., 2014). Then, heritability can be estimated by the fraction of phenotypic variation explained by the genetic relationships among these related subjects. With genome-wide genotyping data on unrelated individuals, an alternative estimator of narrow-sense heritability derives from the additive effects of all common SNPs on phenotype among these unrelated samples, which is usually called SNP heritability (Speed et al., 2016). Instead of using the expected relationship based on pedigree information, SNP heritability is estimated from a genome-wide average across all common SNPs (Toro et al., 2015). Since SNP heritability can capture neither non-additive genetic variation nor genetic variation not covered by SNPs measured by the selected genotyping microarray, it is usually viewed as a lower bound estimate for (narrow-sense) heritability. Recently, computing tools such as genome-wide complex trait analysis (GCTA,Yang et al. (2011)), linkage disequilibrium score regression (Bulik-Sullivan et al., 2015), BOLT-REML (Loh et al., 2015), and massively expedited genome-wide heritability analysis (Ge et al. (2015)) have been developed for SNP heritability estimation.

Heritability is not a fixed property of a phenotype, and analysis of different datasets can result in different estimates of heritability. The estimation of heritability depends on the relative contribution of genetic factors, non-genetic factors and possibly their interaction. People from different ethnic groups can have different genetic backgrounds and be subject to different non-genetic factors. Moreover, methodological factors, such as the sample size of the study and reliability of the phenotype measurement, can also impact the estimation. For these reasons, the United Kingdom (UK) Biobank (Sudlow et al., 2015; Satizabal et al., 2017) provides a unique opportunity to comprehensively study the genetic contributions to many brain phenotypes in one single large-scale, relatively homogeneous population. It is an open-access, large prospective study with over 500,000 participants of middle or elderly age. Around 10,000 of these subjects have brain imaging data available.

Here, we used all common (minor allele frequency [MAF] > 0.01) autosomal SNPs to estimate the heritability for 101 regional brain volumes, including the total brain volume (BV), total grey matter (GM), white matter (WM) and cerebrospinal fluid (CSF). We partitioned genetic variation into individual chromosomes to examine the distribution of heritability across the genome. To assess whether functional annotation (Hu et al., 2017a,b) is associated with genetic effects, we partitioned genetic variation according to cell-type-specific annotations. In addition, we estimated the heritability of principal components (PCs) derived from the regional volume data and evaluated the variability of heritability estimations across brain regions and functional networks. Furthermore, we estimated gender-specific heritability in each region. We compared the findings from the UK Biobank with those from the Alzheimers Disease Neuroimaging Initiative (ADNI, Weiner et al. (2013); *n* ~ 1100), Philadelphia Neurodevelopmental Cohort (PNC, Satterthwaite et al. (2014); *n* ~ 600), and Pediatric Imaging, Neurocognition, and Genetics (PING, Jernigan et al. (2016); *n* ~ 500), which demonstrated that more stable estimates can be obtained from the UK Biobank.

## RESULTS

### Heritability estimates by all common autosomal SNPs

We first estimated the proportion of variation in regional brain volumes that can be explained by all common autosomal SNPs, using linear mixed-effect models (LMMs, see Section Online Methods). Genetic similarity among individuals was captured by the genetic relationship matrix (GRM). We used GCTA tools (Yang et al., 2011) for heritability estimation, adjusting for baseline age, gender, top 10 PCs, as well as BV (to remove scaling effects for other regions).

**Supplementary Tables 1 and 2** display the heritability estimates, standard errors, and p-values from the one-sided likelihood ratio test in each brain region. We found that a large proportion of variation in regional volume is explained by additive genetic effects. The heritability estimates vary across the brain (**Fig. 1**). The top 10 regions with high heritability estimates are the brain stem (82.7%), cerebellar vermal lobules VIII.X (68.3%), cerebellar vermal lobules I.V (68.0%), BV (65.9%), left cerebellum exterior (64.1%), right cerebellum exterior (63.2%), WM (62.8%), right ventral diencephalon (DC) (62.4%), left ventral DC (58.8%), and right cerebellum WM (58.1%), in descending order of heritability point estimate. Noticeable evidence of symmetry in heritability estimates is observed in many brain regions. In Figure 1, many left/right pairs of regions (such as R/L 07, R/L 44, R/L 08, R/L 19) are located next to each other. Since we have a sufficiently large sample size, p-values for most regions are highly significant even after controlling the false discovery rate at 0.05 by Benjamini and Hochberg (1995) procedure.

**Figure 1:**
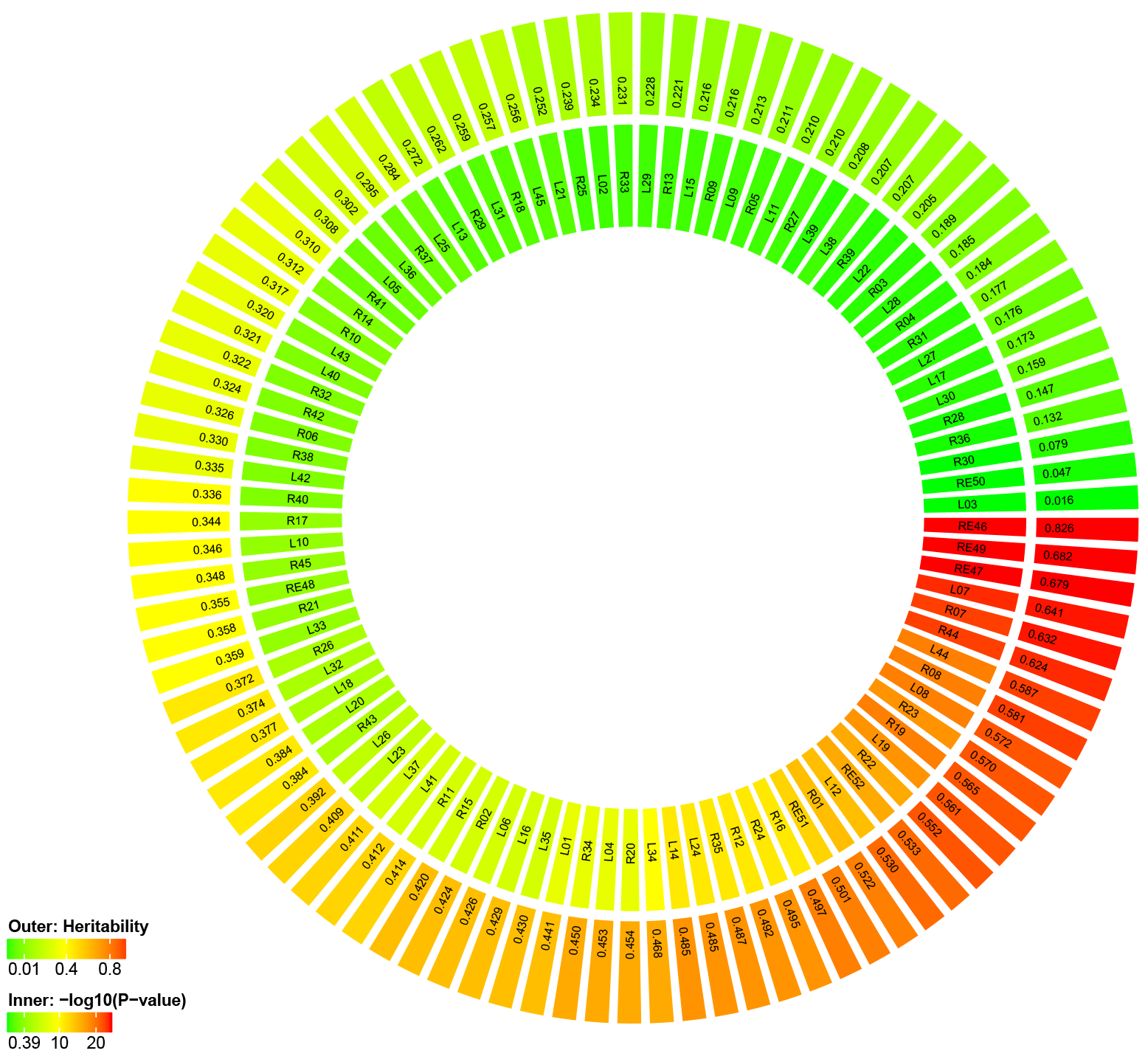
UK Biobank, SNP heritability and adjusted p-values ranked by estimates

We investigated whether the observed considerable variability in heritability estimates across brain regions is due to varying levels of reliability underlying the measurements of these regional brain volumes. **Supplementary Figure 1(a)** shows the relationship between the SNP heritability estimate and the average volume of each brain region, with the latter as a metric to gauge the level of measurement reliability underlying regional brain volumes. While two regions with low heritability estimates also have low average volume size, we observe that high reliability does not necessarily lead to high heritability estimates. Genetic contributions are different among regions with comparable average volume sizes.

We also estimated gender-specific heritability in each region (**Fig. 2**). The top regions with largest gender disparity, as measured by absolute difference in point heritability estimates are listed in **Supplementary Table 3**. Although there are several regions showing strong evidence of gender difference (such as right/left putamen), the distribution of heritability is largely consistent among all, female and male subjects.

**Figure 2:**
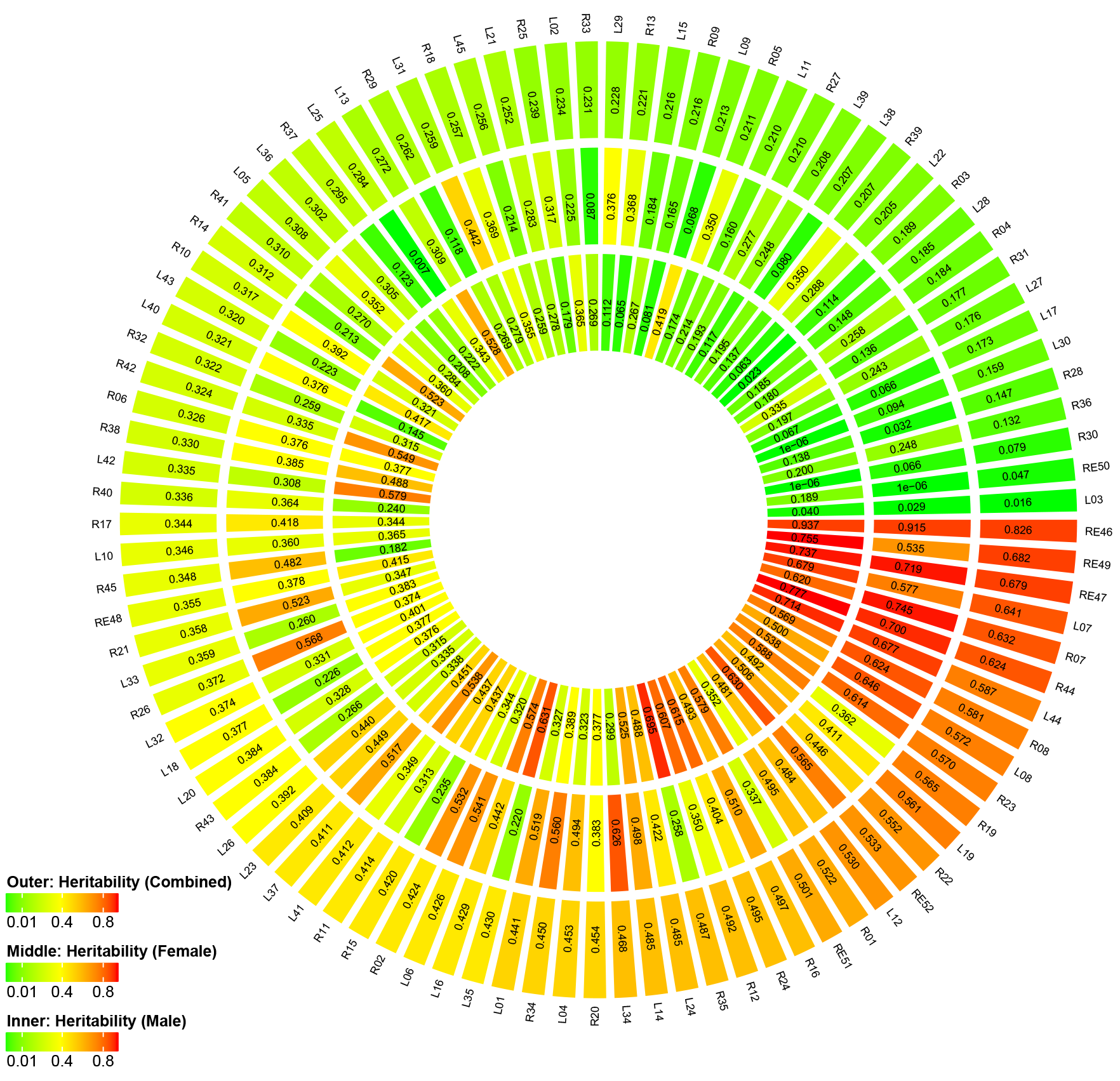
Gender-specific heritability estimate in each region

### Partitioning genetic variation by chromosome

To examine the distribution of heritability across the genome, we partitioned genetic variation into individual chromosomes. Specifically, we estimated GRM using SNPs on each chromosome and estimated heritability separately for each chromosome on each regional brain volume (22 analyses per region, 2222 analyses in total).

**Supplementary Figure 2(a)** shows the heritability estimates by chromosome. The chromosomes are ordered from left to right by their lengths. We found that longer chromosomes tend to have larger heritability estimates than shorter ones. We then calculated the aggregate heritability across all of the 101 regions and found that the aggregated heritability explained by each chromosome is also highly correlated with chromosome length (**Fig. 3(a)**, *R*^2^ = 69.0%, p-value=1.67 × 10^−06^). These findings are consistent with a highly polygenic, or omni-genic model (Lee et al., 2012; Boyle et al., 2017) and indicate that SNPs contributing to variations in regional brain volumes are spread nearly uniformly across the genome.

**Figure 3:**
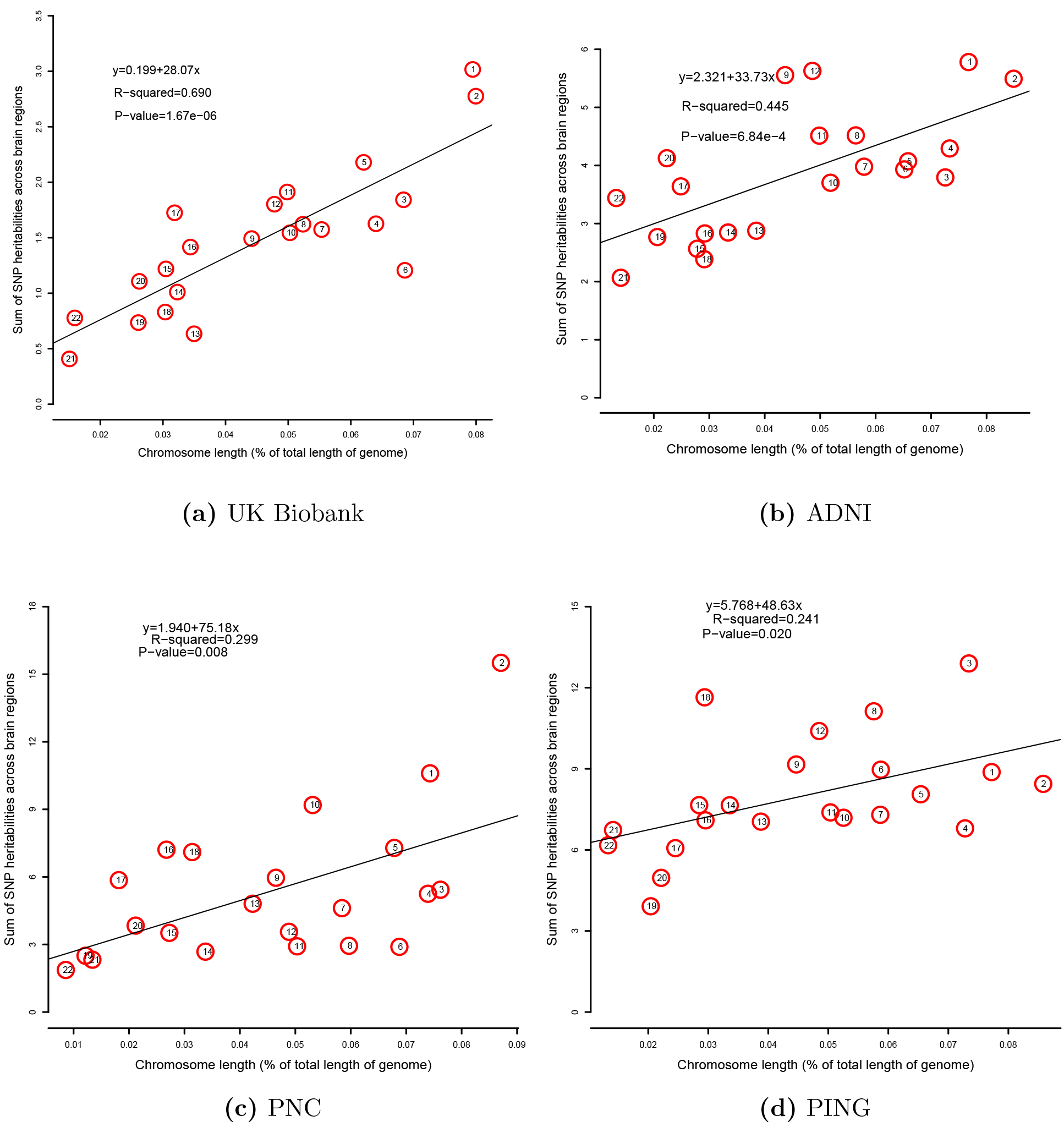
Aggregated heritability of brain regions by each chromosome. In each dataet, heritability explained by each chromosome is highly correlated with chromosome length.

### Partitioning genetic variation by functional annotation

We explored whether functional annotation of SNPs can explain the amount of genetic variation. Following Finucane et al. (2015), we used 220 cell-type-specific annotations. Specifically, SNPs were divided into seven groups according to their activeness among 10 cell groups, namely adrenal gland and pancreas, central nervous system (CNS), cardiovascular system, connective tissue and bone, gastrointestinal, immune and hematopoietic systems, kidney, liver, skeletal muscle and other. In our analysis, we particularly focused on SNPs active in the CNS cell group (see Section Online Methods). We found strong evidence that SNPs residing in chromatin regions inactive across all cell groups contributed less to heritability than SNPs residing in chromatin regions active in at least one cell group. Moreover, SNPs in chromatin regions particularly active in the CNS cell group contributed slightly more to heritability than SNPs in chromatin regions inactive in the CNS cell group (but active in other cell groups). On average, SNPs residing in chromatin regions active in both the CNS cell group and broadly active in other cell groups explain most of the variation in regional volumes (**Fig. 4**).

**Figure 4:**
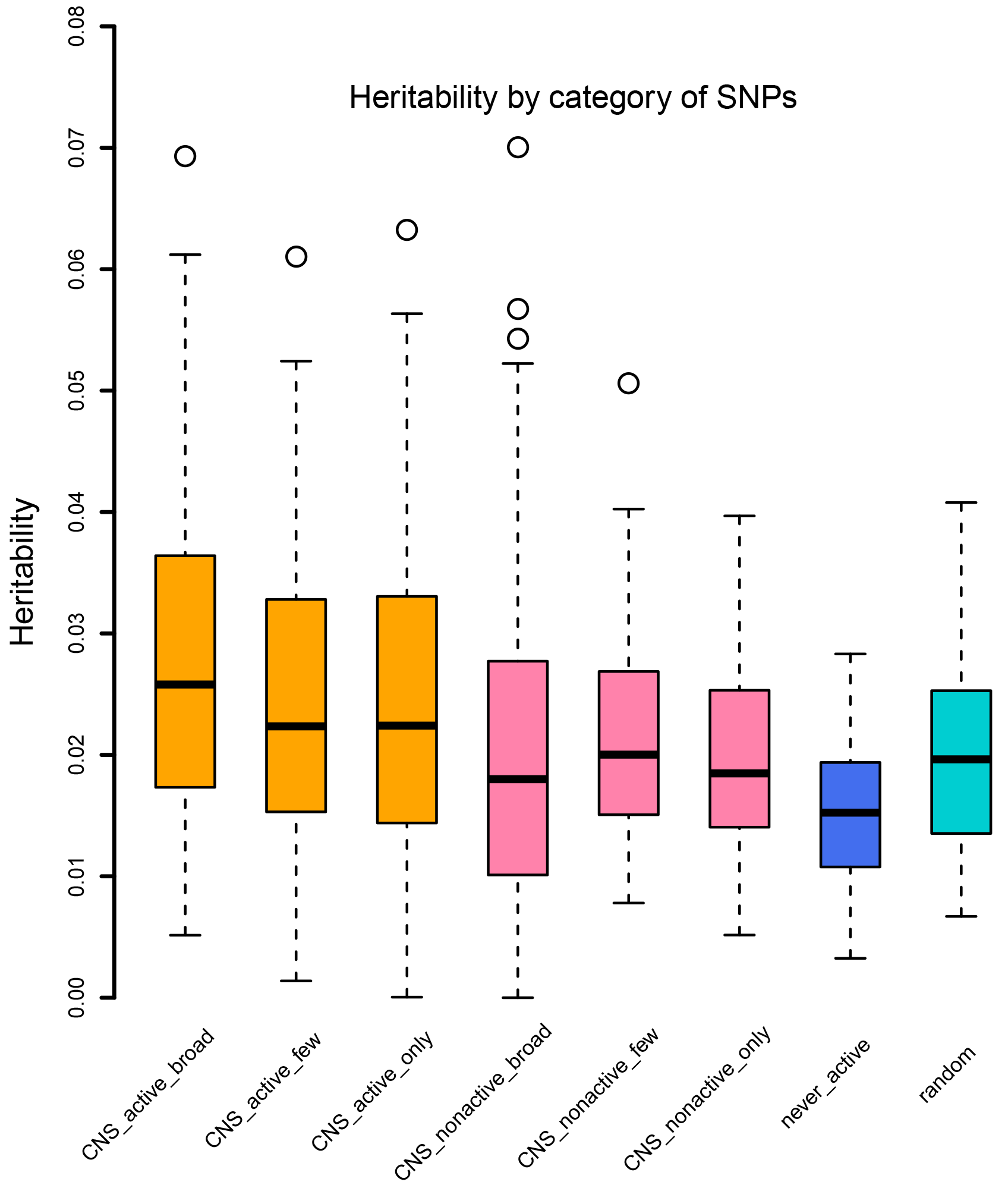
Heritability of brain regions by category of SNPs according to functional annotations

### Heritability pattern across brain function networks

To investigate the heritability pattern across large-scale brain functional networks, we clustered 97 brain regions into 18 functional communities (Buckner et al., 2008; Sporns and Betzel, 2016; Huang et al., 2017). We found that the heritability estimates vary substantially across these functional communities, while the degree of gene control on these functional communities is comparable (**Fig. 5**). Communities with complex functions tend to have large regional variance in heritability. For example, communities C1 and C5 are involved in several networks, including default mode, somatomotor, visual, attention, and language. Regions within the two communities have large variance in heritability estimates. Other clusters linked to simpler functions (with smaller cluster size as well) tend to have smaller regional variance in heritability estimates. The heritability estimates cluster rather tightly together for regions within communities C9 (default mode, motion), C11 (visual), C13 (auditory, language), C14 (memory) and C15 (somatosensory).

**Figure 5:**
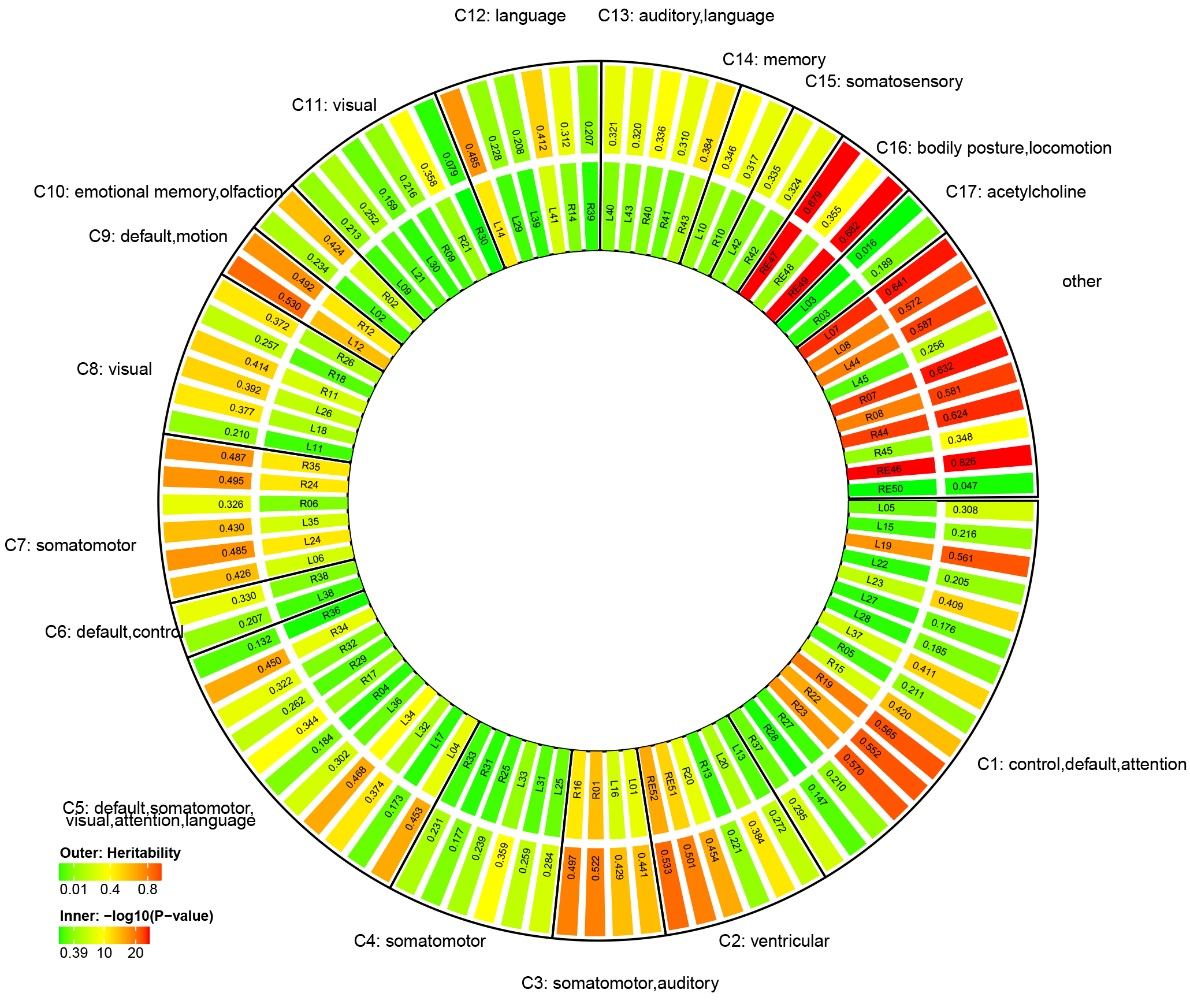
UK Biobank, SNP heritability and adjusted p-values grouped by brain function networks

### Heritability analysis after dimension reduction

We performed principal component analysis (PCA) on the regional brain volumes and obtained the top 10 PCs. **Supplementary Table 4** lists the heritability estimates for the top 10 PCs with and without adjusting for BV. We found that the first PC has a high heritability estimate without adjusting for BV (68.7%), but the heritability estimate is zero after adjusting for BV. These estimates indicate that the first PC fully captured the variance of BV. The Pearson correlation between the first PC and BV is 0.979. As the PCs are orthogonal, adjusting for BV did not affect the heritability estimates of other PCs.

Although the top 10 PCs are highly heritable, the amount of phenotypic variation explained by each PC does not seem to be related to the heritability of the PC. For example, the heritability of the second PC was much smaller than that of the other top 10 components. This result may indicate that although the gene had a large influence on the brain volumes, these phenotypes were not fully genetically controlled by all SNPs in this population. Non-genetic factors, non-additive genetic effects and even batch effects may also contribute to variation in brain volumes. We also calculated heritability estimates by each chromosome (**Supplementary Fig. 3(a)**) for these top 10 PCs and found that the sum of the heritability values explained by each chromosome is again highly correlated with chromosome length (**Supplementary Fig. 4(a)**, *R*^2^ = 49.6%, p-value=2.48 × 10^−04^).

Similar to brain functional community analysis, we grouped the brain regions into 10 modules according to their loadings for the top 10 PCs. That is, we classified the regions corresponding to the top 10 loadings of each component into one module. Each region therefore can fall into more than one module. In our analysis, most regions fell only into one (44 regions) or two (25 regions) modules. **Supplementary Figure 5** shows the distribution of heritability estimates across these 10 modules. Again, regions classified in modules corresponding to PCs that explain more volume variation do not necessarily have higher heritability estimates. As expected, regions in modules corresponding to PCs with higher heritability estimates also have higher heritability estimates.

### Comparing UK Biobank results with results from other datasets

The same analyses presented above in the UK Biobank were conducted in three other datasets, namely ADNI, PNC and PING datasets. Due to smaller sample sizes or less reliable brain imaging data, heritability estimates from these three datasets have much larger variance than those from the UK Biobank (**Fig. 6**, **Supplementary Figs. 6 and 7**). After multiple testing adjustment, we found few regions or PCs to be significant at a false discovery rate of 0.05 in the three studies (**Supplementary Tables 5-7**).

**Figure 6:**
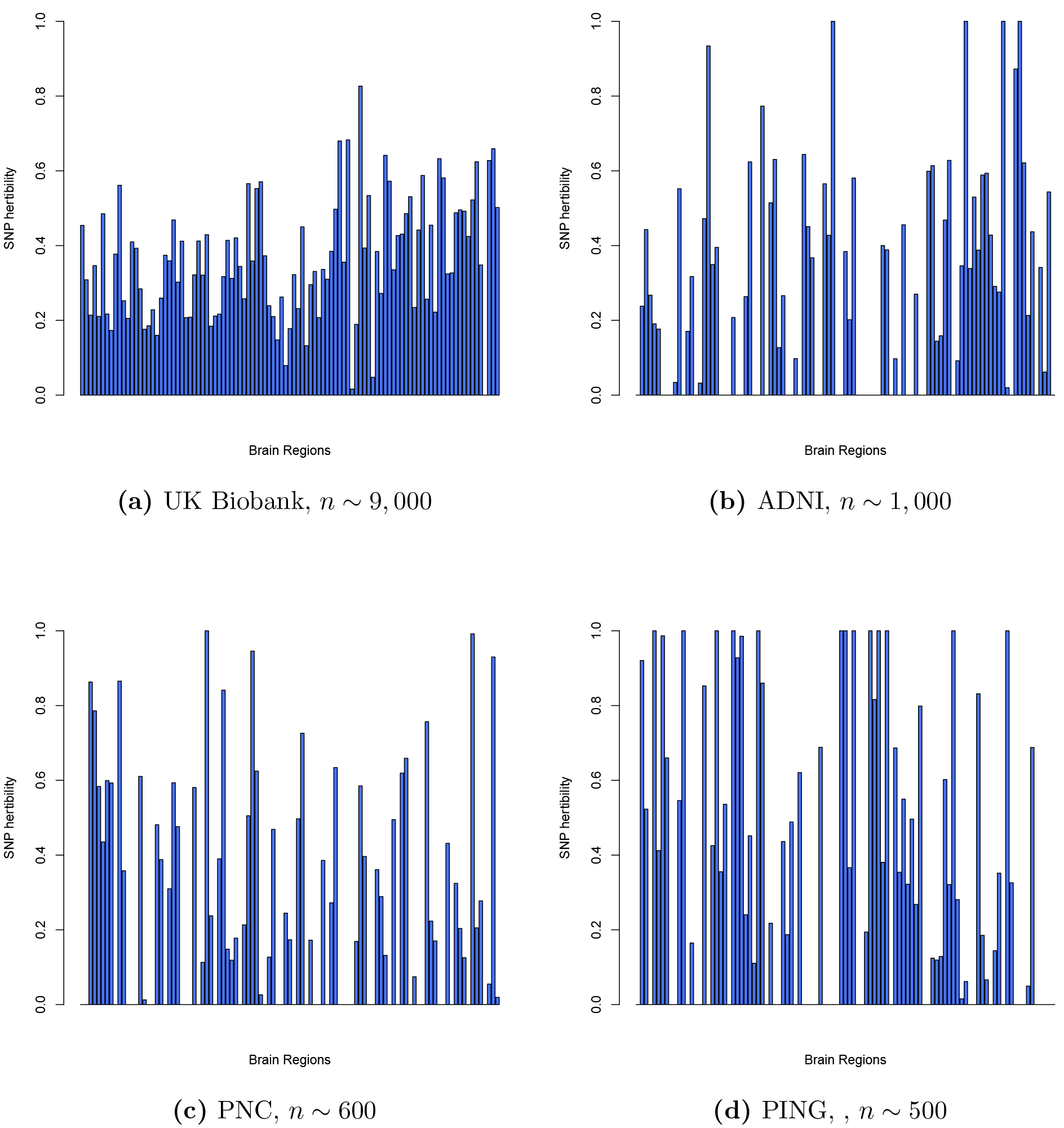
Comparing SNP heritability in different datasets. Estimates of variation explained by all autosomal SNPs of each regional brain volumes as well as GV, WM, BV and CSF (last four bars).

However, some findings are indeed consistent. For example, from each dataset, we observed the linear relationship between chromosome length and the variance explained by each chromosome. But the association tends to be weaker as the sample size decreases (**Fig. 3** and **Supplementary Fig. 4**). In addition, heritability estimates of regional brain volumes are not related to their reliability (**Supplementary Fig. 1**).

## DISCUSSION

In summary, our extensive analyses across four imaging genetic datasets support the following five important findings. First, regional volumes are generally heritable. The majority of brain regions are similarly heritable among females and among males. Study samples used in this work vary from young (PING, PNC) to middle-age/elderly participants (ADNI, UK Biobank). Second, we observe omni-genic patterns where genetic variants contributing to variations in brain volumes are widely spread across the genome with one major evidence being the significant positive linear relationship between chromosome-specific heritability estimates and chromosome length. Third, we found that genetic variants residing in active chromatin regions, particularly those active specifically in the CNS cell group, tend to explain more variation in brain volumes.

Fourth, through PCA, we demonstrated that the top PCs are also highly heritable, but the amount of brain volume variation explained by the PCs does not seem to be related to the heritability estimates of these PCs. Fifth, the genetic influences are not uniformly distributed across the brain regions or brain functional communities. Similar genetic control can be found among regions within a small community and on pairs of regions in the left and right hemispheres. Results from the four independent cohorts are largely consistent. Compared to ADNI, PNC and PING, UK Biobank can provide more stable estimates of heritability with smaller standard errors.

We found that 65.9% of BV variability can be explained by genetic variation of all common autosomal SNPs for UK Biobank subjects. Adjusting for BV, 39.3% of CSF volume variability is explained by genetic variation. Without BV adjustment, the heritability estimates for GM (67.5%) are similar to the estimates for BV (65.9%); after adjusting for BV, the heritability estimates for GM are zero, suggesting that BV and GM share the same or very similar genetic bases. For WM, however, heritability estimates remain almost unchanged before and after adjusting for BV (62.8% and 62.0%, respectively), indicating that genes underlying WM are not general brain growth genes, but rather more likely to be genes that specific control this particular brain structure and sub-regions. Our heritability estimates are similar to those reported in (Pol et al., 2006; Kremen et al., 2010; Carmelli et al., 1998; Bryant et al., 2013). We have more clearly illustrated the different genetic bases behind BV/GM volume and WM volume.

In regional volume analysis, we obtained the heritability estimates of 97 regions, showing that the regions are highly heritable and genetic influences are not uniformly distributed across the brain. To assess whether the lower heritability is caused by the difficulty in accurately measuring the regional volume, we quantify the concordance between the average volume sizes and heritability estimates. We found no evidence that the higher heritability is driven by the higher reliability of the volume measurement. Regional variation in terms of genetic contribution is observed among the regions with comparable average volume sizes. Thus, prioritizing regions with high heritability for genetic studies are more likely to result in reproducible *bona fide* findings. The results are consistent in all four datasets and agree with findings from other studies on brain shape measurements and hippocampal sub-region volumes (Roshchupkin et al., 2016; Whelan et al., 2016). In addition, we found strong evidence that the estimates have a symmetric pattern across the left and right hemispheres. Many left/right pairs of regions have similar estimates, consistent with results from previous twin studies (Chen et al., 2012; Wright et al., 2002). Although several regions have large gender differences in heritability, our gender-specific analysis show that the majority of additive genetic effects are shared between female and male subjects.

To further study the patterns of regional variations in heritability estimates, we clustered the regions by their biological functions. In brain functional network analysis, we grouped the 97 regions into 18 non-overlapping brain functional communities (details can be found in the **Supplementary Note**). We found the community-wise variability in heritability across these functional communities, while the genetic influences widely prevail across the brain functional networks with comparable degrees of control (heritability). The regions within each community do not necessarily have similar heritability estimates, depending on the complexity of the community functions. We performed PCA and found that the components explaining more volume variations do not necessarily have higher heritability, nor higher loadings on regions with higher heritability. This makes sense because PCA is an unsupervised dimension reduction technique. Non-genetic factors or non-additive genetic effects that are not captured by SNPs also influence variation in brain volume.

The significant linear correlation between the variance explained by a chromosome and the length of the chromosome was observed on both the volumes and principal components. These patterns suggest that genetic variants controlling regional brain volumes are rather ubiquitously distributed across the genome. Similar findings have been reported on other phenotypes, such as height, body mass index, neuroanatomical phenotypes and schizophrenia (Yang et al., 2011; Lee et al., 2012; Toro et al., 2015; Fritsche et al., 2016; Shi et al., 2016; Shan et al., 2017; Kemp et al., 2017). To explain this phenomenon, Boyle et al. (2017) proposed an omnigenic hypothesis where most heritability can be explained by effects of genes outside core pathways because gene regulatory networks are sufficiently interconnected. Although SNPs influencing regional brain volumes spread widely across the genome, effect signals are associated with cell-type-specific annotations. For regional brain volumes, we show enrichment of genetic signals in active chromatin regions, especially those that are active specifically in the CNS cell type and broadly active in other cell types.

Finally, we compared the results from UK Biobank with the results from the other three datasets. The UK Biobank allows more stable estimation of the magnitude of genetic determination of the human brain. In ADNI, PNC and PING, extreme estimates such as 0.9999 or 0 occurred for some regions (**Fig. 6**); these estimates should not be interpreted as 'true’ heritability estimates, but only indicate large or small heritability values for a region. Such extreme estimates may be due to insufficient sample size or low reliability of volume measurements. In UK Biobank, no such extreme estimates are observed, and the heritability estimates range from 1.6% to 82.6%, with standard error approximately 0.07. Although SNP heritability estimates are at the lower bound of (narrow-sense) heritability, we observed many heritable brain regions using the UK Biobank dataset, and the estimates are statistically significant using a likelihood ratio test after multiple testing adjustment Benjamini and Hochberg (1995). For the other three datasets, however, few significant findings remain after multiple testing adjustment.

## METHODS

### Methods are available in the Online Methods section

*Note: One supplementary information pdf file and one supplementary Excel file are available.*

## ACKNOWLEDGEMENTS

This research was partially supported by U.S. NIH grants MH086633 and MH092335, NSF grants SES-1357666 and DMS-1407655, a grant from the Cancer Prevention Research Institute of Texas, and the endowed Bao-Shan Jing Professorship in Diagnostic Imaging. We thank the UK Biobank, ADNI, PING and PNC participants for their participation and the research teams for their work in collecting, processing and disseminating these datasets for analysis. This research has been conducted using the UK Biobank Resource (application number 22783), subject to a data transfer agreement. Part of data collection and sharing for this project was funded by the Alzheimers Disease Neuroimaging initiative (ADNI) (National Institutes of Health Grant U01 AG024904) and DOD ADNI (Department of Defense award number W81XWH-12-2-0012). ADNI is funded by the National Institute on Aging, the National Institute of Biomedical Imaging and Bioengineering and through generous contributions from the following: Alzheimers Association; Alzheimers Drug Discovery Foundation; Araclon Biotech; BioClinica, Inc.; Biogen Idec Inc.; Bristol-Myers Squibb Company; Eisai Inc.; Elan Pharmaceuticals, Inc.; Eli Lilly and Company; EuroImmun; F. Hoffmann-La Roche Ltd and its affiliated company Genentech, Inc.; Fujirebio; GE Healthcare; IXICO Ltd; Janssen Alzheimer Immunotherapy Research & Development, LLC; Johnson & Johnson Pharmaceutical Research & Development LLC; Medpace, Inc.; Merck & Co., Inc.; Meso Scale Diagnostics, LLC; NeuroRx Research; Neurotrack Technologies; Novartis Pharmaceuticals Corporation; Pfizer Inc.; Piramal Imaging; Servier; Synarc Inc.; and Takeda Pharmaceutical Company. The Canadian Institutes of Health Research is providing funds to support ADNI clinical sites in Canada. Private sector contributions are facilitated by the Foundation for the National Institutes of Health (www.fnih.org). The grantee organization is the Northern California Institute for Research and Education, and the study is coordinated by the Alzheimers Disease Cooperative Study at the University of California, San Diego. ADNI data are disseminated by the Laboratory for Neuro Imaging at the University of Southern California. Part of data collection and sharing for this project was funded by the Pediatric Imaging, Neurocognition and Genetics Study (PING) (National Institutes of Health Grant RC2DA029475). PING is funded by the National Institute on Drug Abuse and the Eunice Kennedy Shriver National Institute of Child Health & Human Development. PING data are disseminated by the PING Coordinating Center at the Center for Human Development, University of California, San Diego. Support for the collection of the PNC data sets was provided by grant RC2MH089983 awarded to Raquel Gur and RC2MH089924 awarded to Hakon Hakonarson. All PNC subjects were recruited through the Center for Applied Genomics at The Children's Hospital in Philadelphia.

## AUTHOR CONTRIBUTIONS

H.Z., P.L., J.I. and Y.L. designed the study. B.Z. performed the experiment and analyzed the data. T.L., J.Z., Y.W., Y.S., Z.Z., F.Z., C.H., H.L. and J.Y. downloaded the datasets, preprocessed MRI data, undertook the quantity controls and imputed SNP data. B.Z., Y.L., P.L., J.I. and H.Z. wrote the manuscript with feedback from all authors.

## COMPETETING FINANCIAL INTERESTS

The authors declare no competing financial interests.

## ONLINE METHODS

### Participants and image preprocessing

Datasets used in this paper included the UK Biobank, ADNI, PNC, and PING. Detailed data collection/processing procedures and quality control prior to the release of data are documented at http://www.ukbiobank.ac.uk/resources/ for UK Biobank, http://adni.loni.usc.edu/data-samples/ for ADNI, http://pingstudy.ucsd.edu/resources/genomics-core.html for PING and https://www.ncbi.nlm.nih.gov/projects/gap/cgi-bin/study.cgi?study_id=phs000607.v1.p1 for PNC. For each dataset, we used subjects with both magnetic resonance imaging (MRI) and SNP data available after applying proper quality controls. We only used baseline data for longitudinal studies.

The MRI data were preprocessed using standard procedures via advanced normalization tools (ANTs, Avants et al. (2011)). Following Avants et al. (2011)), our preprocessing steps consisted of the N4 bias correction, registration-based brain extraction, and a prior-based N4-Atropos 6 tissue segmentation (oasis template), which classified the brain into WM, GM, deep GM, CSF, brainstem and cerebellum. We then adopted the 101 regions of interest (ROIs) defined by the manually edited labels of the publicly available MindBoggle-101 dataset (Klein and Tourville, 2012) to perform multi-atlas cortical parcellation.

We excluded subjects for whom the imaging data did not pass the standard imaging quality controls, and removed three ROIs with many missing values: X5th ventricle, left lesion and right lesion. There was a total of 101 regional brain volumes, including total BV, GM, WM and CSF. We standardized each volume to better fit the assumption for the LMM. By checking the studentized residuals of the LMM between volume with age and gender, we deleted the top 10 outlier subjects for each standardized volume. The demographic information related to the MRI datasets are listed in Supplementary Table 11.

### Genotyping

Genotype imputation was performed on the PNC, ADNI, and PING datasets. For UK Biobank, we used an unimputed dataset. Standard quality controls were performed to ensure high quality of the SNP data. These procedures were performed using the Plink tool set (Purcell et al., 2007).

### PNC

From the PNC database, 8722 participants were genotyped on one of the six different platforms: 66 were genotyped on the Affymetrix array 6.0; 722 were genotyped on the Axiom array; 556 were genotyped on the Illumina HumanHap 550 array version1; 1914 were genotyped on the Illumina HumanHap 550 array version 3; 1657 were genotyped on the Illumina HumanHap 610 array; and 3807 were genotyped on the Illumina Human Omni Express array. We applied the quality control steps to each dataset separately, which included removal of subjects with more than 10% missing values, removal of SNPs (i) with more than 5% missing values, (ii) with MAF smaller than 5%, (iii) with Hardy-Weinberg equilibrium test p-value < 1 × 10^−6^, and (iv) located on a sex chromosome. We then employed MACH-Admix software (Liu et al., 2013) to perform genotype imputation, using 1000G Phase I Integrated Release Version 3 haplotypes (1000-Genomes-Project-Consortium et al., 2012) as a reference panel. We also conducted quality control after imputation, excluding markers with (i) low imputation accuracy (based on imputation output *R*^2^); and (ii) Hardy-Weinberg equilibrium test p-value < 1 × 10^−6^. We combined the six datasets and retained the shared SNPs. Finally, 5, 354,265 bi-allelic markers (including SNPs and indels) from 8681 subjects remained for further analysis.

### ADNI

Genetic data from ADNI1 (620,901 genetic markers from 818 subjects) and ADNI2/GO (730, 525 genetic markers from 432 subjects) were processed separately with the following pipeline. The first-line quality control steps include (i) call rate check per subject and per SNP marker, (ii) gender check, (iii) sibling pair identification, and (iv) population stratification. The second-line preprocessing steps include removal of SNPs (i) with more than 5% missing values, (ii) with MAF smaller than 10%, (iii) with Hardy-Weinberg equilibrium p-value < 1 × 10^−6^ and (iv) located on a sex chromosome. For further processing, we included 503, 778 SNPs from 756 white (Caucasian) subjects from ADNI1 and 516,453 SNPs from 397 white subjects from ADNI2/GO. We employed MACH-Admix software (Liu et al., 2013) to perform genotype imputation, using 1000G Phase I Integrated Release Version 3 haplotypes (1000-Genomes-Project-Consortium et al., 2012) as a reference panel. We conducted quality control after imputation, excluding markers with (i) low imputation accuracy (based on imputation output *R*^2^); and (ii) Hardy-Weinberg equilibrium p-value < 1 × 10^−6^. We then had 7, 986, 566 bi-allelic markers (including SNPs and indels) from 756 subjects from ADNI1 and 8,218,182 markers from 397 subjects from ADNI2/GO. We combined the two datasets and retained the shared SNPs. Finally, 7, 664, 643 bi-allelic markers (including SNPs and indels) from 1153 subjects remained for further analysis.

### PING

We applied the following preprocessing technique to the genetic data. The first-line quality control steps included (i) call rate check per subject and per SNP marker, (ii) gender check, and (iii) sibling pair identification. The second-line preprocessing steps included removal of SNPs (i) with more than 5% missing values, (ii) with MAF smaller than 10%, (iii) with Hardy-Weinberg equilibrium p-value < 1 × 10^−6^, and (iv) located on a sex chromosome. We thus had 539, 865 SNPs from 1036 subjects for further processing. We employed MACH-Admix software (Liu et al., 2013) to perform genotype imputation, using 1000G Phase I Integrated Release Version 3 haplotypes (1000-Genomes-Project-Consortium et al., 2012) as a reference panel. We also conducted quality control after imputation, excluding markers with (i) low imputation accuracy (based on imputation output *R*^2^), and (ii) Hardy-Weinberg equilibrium p-value < 1 × 10^−6^. Finally, 10,883, 584 bi-allelic markers (including SNPs and indels) from 1036 subjects were retained for data analysis.

### Further quality control

On each SNP dataset, we further selected subjects with available brain volume data. We then used all autosomal SNPs and again applied the standard quality control procedures: excluding subjects with more than 10% missing genotypes, only including SNPs with MAF > 0.01, with genotyping rate > 90%, and passing Hardy-Weinberg test (*P* > 1 × 10^−7^). We further removed non-European subjects, if any. In PING, we only used biologically unrelated subjects. After quality control, we calculated the GRM by all SNPs and by SNPs on each chromosome separately using GCTA software (Yang et al., 2011). To avoid including closely related relatives, we excluded one of any pair of individuals with estimated genetic relationship larger than 0.025. The sample sizes of the datasets after conducting all quality control procedures are listed in **Supplementary Table 12**.

### Heritability analysis

First, for each regional volume, we estimated the proportion of variation explained by all autosomal SNPs with a LMM (101 analyses in total). The formal setting of the LMM and definition of likelihood ratio test statistics can be found in Yang et al. (2011). The basic idea is to fit the GRM with random effects to the phenotypic measure, while adjusting for other covariates with fixed effects. The GRM was the correlation matrix of participants estimated by the common genetic variants, which was expected to capture the genetic similarity among unrelated individuals. Then the heritability of a phenotype was estimated by contrasting the genetic similarity among individuals with their phenotypic similarity. Baseline age, gender indicator, top 10 PCs of GRM, and BV (for regions other than BV itself) were included as covariates, unless otherwise stated. We also included the phase indicator for the ADNI study to adjust for potential batch effects. Besides the combined sample, we fitted the LMM separately on female and male samples for UK Biobank data.

Second, we partitioned the genetic variation by each chromosome. We estimated the GRM of each chromosome and fitted each of them separately on each volume (22 analyses per volume, 2222 analyses in total). The same set of covariates was included in these LMMs.

Next, we performed PCA on the volumes and computed the heritability of the top 10 PCs. We also partitioned the genetic variation on the components by each chromosome. In the LMMs for the components, we did not adjust for BV unless otherwise stated, since we have observed that the variation of BV is almost captured by the first component, and should be orthogonal to the remaining components.

Finally, we fitted linear models between the length of a chromosome and the aggre-gate heritability of all volumes or their components to study the heritability distribution across the genome. We clustered the regions according to their biological functions and showed the heritability distribution across these communities using the R package circlize (Gu et al., 2014).

### Functional enrichment of genetic signals

Cell-type-specific active chromatin annotations per SNP were from Finucane et al. (2015) and Boyle et al. (2017) (https://github.com/bulik/ldsc/wiki/Partitioned-Heritability). According to Finucane et al. (2015), we performed functional annotation analyses using cell-type-specific annotations marked by the four histones: H3K4me1, H3K4me3, H3K9ac and H3K27ac. Each cell-type-specific annotation corresponded to a histone mark in a single cell type, and there were 220 such annotations. The 220 cell-type-specific annotations were further divided into 10 groups, including adrenal gland and pancreas, CNS, cardiovascular system, connective tissue and bone, gastrointestinal, immune and hematopoietic systems, kidney, liver, skeletal muscle and other. The SNPs were first divided into four overlapping groups according to their activeness in all cell-type groups (only, few, broad, and never active). A SNP was labeled ‘only’ if it was annotated as active in only one of the 10 cell-type groups. A SNP was labeled ‘few’ if it was annotated as active in at most 5 cell-type groups. SNPs that were active in 6−10 cell-type groups were labeled ‘broad’, and SNPs that were not active in any cell type were labeled ‘never active’. Then, SNPs were further labeled as either active in the CNS cell group (‘CNS active’) or not (‘CNS inactive’). As the number of SNPs in each group was different, we randomly selected the same number of SNPs from each cell group (n=8368) and computed the heritability for each group in each region. We generated 50 random selected SNP datasets and calculated the mean of these 50 heritability estimates in each region.

### Data availability

Links to all datasets (UK Biobank, ADNI, PNC and PING) that support the findings of this study are provided in Section Online Methods. Researchers can apply to use these datasets for health related research in the public interest.

